# Integrative omics analysis of *Pseudomonas aeruginosa* virus PA5oct highlights the molecular complexity of jumbo phages

**DOI:** 10.1101/679506

**Authors:** Cédric Lood, Katarzyna Danis-Wlodarczyk, Bob G. Blasdel, Ho Bin Jang, Dieter Vandenheuvel, Yves Briers, Jean-Paul Noben, Vera van Noort, Zuzanna Drulis-Kawa, Rob Lavigne

## Abstract

*Pseudomonas* virus vB_PaeM_PA5oct is a compelling model phage to investigate phage-bacteria interactions and is a potent candidate for phage therapy. Combining hybrid genome sequencing, RNA-Seq, and mass spectrometry of virion-associated proteins allowed us to accurately assemble and annotate PA5oct’s 286,783 bp genome. It encodes 461 coding regions, of which 93 were identified as virion-associated proteins using ESI-MS/MS. RNA-seq revealed twelve tRNA genes, four ncRNA regions, and a gradual take-over of the total cell transcriptome from 21% in early infection to 93% in late infection. Although PA5oct is not organized into strictly contiguous regions of temporal transcription, defined genomic regions are expressed in early, middle, and late phases of infection. Interestingly, we observed distinct regions showing limited transcription activity throughout the infection cycle. The infection of *Pseudomonas aeruginosa* by PA5oct triggers the upregulation of bacterial operons related to NAD biosynthesis, Psl exopolysaccharide, and periplasmic nitrate reductase production and downregulates Type IV pili gene products, suggesting mechanisms of superinfection exclusion. We used the proteome of PA5oct to position this isolate amongst other phages using a protein-sharing network. This study expands our understanding of the molecular diversity of jumbo viruses and opens new questions towards cellular regulation and phage-encoded hijacking mechanisms.

## Introduction

The discovery of very large microbial viruses infecting bacteria and protozoa has dramatically expanded the range of known viral genome sizes, starting around 2 kb and ending upward of 2 Mb. These so-called jumbo viruses can have genomes that are larger than those of parasitic bacteria and archaea, thereby bridging the former classical divide between cells and viruses in terms of genome size (Claverie *et al.*, 2009; Koonin *et al.*, 2015). Importantly, these uncharacterized viral genomes are diverse, and most of their genes lack nucleotide similarity to other functionally characterized or indeed even known sequences that can be accessed in public databases. As such, they have been aptly described as “Viral Dark Matter” and there is room for the exploration of this untapped sequence space (Hatfull, 2015).

Bacteriophages with genome sizes beyond 200 kbp are known as jumbo phages, sometimes also referred to as giant phages. They have been isolated from a multitude of environments, including water, soil, plants, and animal tissues (Yuan *et al.*, 2017). A subset of 120 of them have been sequenced to date (May 2019, Supplementary Table S1), including 109 myoviruses, 10 siphoviruses, and 1 unclassified phage. Most infect Gram-negative bacteria (95.4%), with only a few (11 jumbo phages) infecting Gram-positive bacteria, mostly *Bacillus* strains. The tiny plaques typically formed by these large-sized phages are smaller than 0.5 mm on 0.7% soft agar and are easily overlooked during classical propagation procedures. Their large virion size can also lead to their loss during filtration procedures that are standard in many phage isolation protocols (Sewer *et al.*, 2007), suggesting biases against their isolation.

Although jumbo phages have proven to be interesting for structural analysis and have provided insights into infection processes (Fokine *et al.*, 2007; 2005; Wu *et al.*, 2012), the understanding of their functional genetics and genome organization has often remained limited to the results of *in silico* predictions of genomes assembled from short-reads sequencers such as Illumina machines. However, investigation into their basic (molecular) microbiology, including elucidating their transcriptional schemes, or the mechanisms of their co-evolution with their host must be pursued further if we are to acquire new insights (Hendrix, 2009). RNA-sequencing has proven to be an important tool in that respect, allowing a detailed molecular elucidation of phage transcriptional schemes, and the discovery of regulatory elements and sRNAs, but also providing clues towards the host response during phage infection and its evolutionary implications (Ceyssens *et al.*, 2014; Chevallereau *et al.*, 2016; Blasdel *et al.*, 2017b and 2018).

*Pseudomonas* virus phiKZ, the first sequenced jumbo phage (Mesyanzhinov *et al.*, 2002) serves as a model phage in this regard. In particular, its transcription scheme is independent from the host transcriptional machinery and is governed by the consecutive action of phage-encoded, non-canonical multisubunit RNAPs (virion associated, vRNAP), distantly related to bacterial β and β’ subunits, that are co-packaged alongside the genome and co-injected upon infection to bootstrap the transcription of other RNAP genes present in the phage genome (Ceyssens *et al.*, 2014).

Here we present a detailed molecular analysis of the jumbo *Pseudomonas* virus vB_PaeM_PA5oct, a myovirus infecting *Pseudomonas aeruginosa* (Drulis-Kawa *et al.*, 2014) that lacks a phage-encoded RNAP and thus represents a fundamentally different transcription mode from phage phiKZ. Our investigation was driven by combining short and long read sequencing, temporal transcriptomics, and structural proteomics. Together, these molecular techniques allowed to elucidate the genome organization of this phage species, that currently has no close representatives in the public databases in terms of DNA homology, and the response it elicits during infection of its host *P. aeruginosa* PAO1. Furthermore, we establish the evolutionary relationships between PA5oct and other jumbo bacteriophages using a protein-sharing network analysis.

## Material & Methods

### Bacteriophage propagation, purification, and morphology

In all experiments, phage PA5oct was propagated as previously described (Danis-Wlodarczyk *et al.*, 2015). Phage lysate was purified by 0.45 and 0.22 μm filtration and incubation with 10% polyethylene glycol 8000 (PEG 8000)–1 M NaCl according to standard procedures (Ceyssens *et al.*, 2006). Finally, CsCl-gradient ultracentrifugation was applied (Ceyssens *et al.*, 2008) and the resulting phage preparation dialyzed three times for 30 min against 250 volumes of phage buffer using Slide-A-Lyzer Dialysis Cassettes G2 (Thermo Fisher Scientific Inc, MA, USA). The phage titer was assessed using the double-agar layer technique (Adams, 1959) and purified samples were stored at 4°C.

### The rifampicin growth assay

One-step growth curves were established according to the method of Pajunen *et al.* (2000), with modifications. An equal volume of bacterial culture at OD_600_ of 0.4 was mixed with phage suspension (10^6^ pfu/ml) to obtain a MOI of 0.01. Phages could adsorb for 8 min at 37°C, after which the mixture was diluted to 10^−4^. Triplicate samples were taken every 5 min during 1-1.5 h for titration, (Danis-Wlodarczyk *et al.*, 2015). Where appropriate, rifampicin (Sigma) was added to a final concentration of 400 μg/ml.

### DNA isolation and hybrid sequencing

Phage genomic DNA was prepared with the use of a modified protocol for lambda DNA isolation according to Ceyssens (2009) and sequenced using the Illumina MiSeq platform available at the Nucleomics Core (VIB, Belgium) as previously described by Danis-Woldarczyk *et al.* (2015). The phage genomic DNA was also prepared for long reads sequencing using Oxford Nanopore Technology (ONT) 1D library protocol as described in Kakabadze *et al.* (2018). The library was sequenced in-house on a MinION equipped with a R9.4.1 flowcell (ONT).

### RNA extraction and sequencing

*P. aeruginosa* strain PAO1 was grown overnight in 5 ml LB medium at 37 °C. Next, cells were diluted 1:100 in 50 ml fresh medium and further grown at 37 °C until an OD_600_ of 0.3 was reached (around 1.2 × 10^8^ CFU/mL, early exponential phase). The culture was infected with PA5oct at a MOI of 50. To control for a synchronous infection, we ensured that less than 5% of bacterial survivors remained after five minutes post infection. Biologically independent samples were collected in triplicates at 5 minutes (early), 15 minutes (middle), and 25 minutes (late) into infection and processed as described previously (Blasdel *et al.*, 2017a; Chevallereau *et al.*, 2016).

### Genome and transcriptome analysis

The genome of PA5oct was reconstructed using both Nanopore and Illumina reads with the integrated hybrid assembler Unicycler v0.4.7 (Wick *et al.*, 2017). After assembling in a single circular contig, the ends of the genome were determined by mapping the sequencing reads to the genome using bwa-mem v0.7.17 (Li H, 2013), and by analysis of Illumina reads with PhageTerm v0.12 (Garneau *et al.*, 2017). The genome architecture was also confirmed by investigating potential structural variations using the long nanopore reads with a combination of the tools ngmlr and sniffles (Sedlazeck *et al.*, 2018), and the genome browser ribbonviewer (Nattestad *et al.*, 2016).

The genome annotation was performed by integrating the RAST phage annotation tool (McNair *et al.*, 2018) and a manual curation with standard genome annotation methods (Danis-Wlodarczyk *et al.*, 2015; and 2016). The annotation was further inspected visually with UGENE (Okonechnikov *et al.*, 2012) using the stranded RNA-Seq reads, mapped to the phage genome. The RNA-sequencing reads were aligned to both the phage and host genomes using the software TopHat v2 (Kim *et al.*, 2013) and the resulting alignments were then summarized strand-independently into count tables using HTSeq-Count (Anders *et al.*, 2015). Finally, the counts of each features were normalized using TPM (transcripts per million). For the differential expression of the host features, each statistical comparison presented was performed using the DESeq2 (Love et al. 2014) R/Bioconductor package to normalize host transcript populations to host transcript populations, or phage to phage, before testing for differential expression as described by Blasdel *et al.* (2018).

The genome of PA5oct was deposited under the NCBI Genbank accession number MK797984, Illumina and Nanopore genomic reads are available in the NCBI SRA database via the bioproject accession number PRJNA516157, and the transcriptomics dataset is available through the NCBI GEO database via the accession number GSE130190.

### ESI-MS/MS analysis on virion particle proteins

Phage proteins were isolated from a purified phage lysate (10^9 pfu/ml) by a single methanol/chloroform extraction (1:1:0.75, v/v/v) (Acros Organics) and precipitated by addition of an equal volume of methanol (14,000 x *g*, 6 min). The phage proteins were separated on a 12% SDS-PAGE gel and analyzed by ESI-MS/MS as previously described (Ceyssens *et al.*, 2014; Van den Bossche *et al.*, 2014).

### Protein family clustering and construction of the relationships network

To build a gene-sharing network, we retrieved 231,166 protein sequences representing the genomes of 2,304 bacterial and archaeal viruses from NCBI RefSeq (version 85) and used the network analytics tool, vConTACT (version 2.0; https://bitbucket.org/MAVERICLab/vcontact2) (Jang *et al.*, 2019), as an app at iVirus (Bolduc *et al.*, 2017a). Briefly, including protein sequences from PA5oct, a total of 231,627 sequences were subjected to all-to-all BLASTp searches, with an E-value threshold of 10^-4^, and defined as the homologous protein clusters (PCs) in the same manner as previously described (Bolduc *et al.*, 2017b). Based on the number of shared PCs between the genomes, vConTACT v2.0 calculated the degree of similarity as the negative logarithmic score by multiplying hypergeometric similarity P-value by the total number of pairwise comparisons. Subsequently, pairs of closely related genomes with a similarity score of ≥ 1 were grouped into viral clusters (VCs), with default parameters of vConTACT v2.0 (Jang *et al.*, 2019). The network was visualized with Cytoscape (version 3.5.1; http://cytoscape.org/), using an edge-weighted spring embedded model, which places the genomes or fragments sharing more PCs closer to each other. The taxonomic affiliation was taken from the International Committee on Taxonomy of Viruses (ICTV Master Species List v1.3; http://www.ictvonline.org/virusTaxonomy) incorporated 2017 ICTV updates (Adams *et al.*, 2017; Adriaenssens *et al.*, 2017). Further, the proportion of shared PCs between two genomes was characterized, as previously described (Bolduc *et al.*, 2017b and Jang *et al.*, 2019).

## Results

### Basic microbiological and genome properties of *Pseudomonas* phage PA5oct

Under standard conditions, one-step growth experiments indicate a latent period of approximately 40 min and a burst size of about 30 - 40 phage particles per infected bacterial cell (Drulis-Kawa *et al.*, 2014). When 400 μg/ml of rifampicin, a host RNA polymerase (RNAP) transcription inhibitor, is added prior PA5oct infection, the progeny phage production is completely suppressed. This suggests that PA5oct, in contrast to another jumbo phage *Pseudomonas* virus phiKZ, relies on the host RNAP for phage transcription.

Phage PA5oct has a linear, A+T-rich (33.3% GC-content), double-stranded DNA genome of 286,783 bp with 461 putative CDS that could be validated by inspection with the RNA-seq data (Figure 1). In comparison, the *in silico* RAST annotation pipeline (McNair *et al.*, 2018) allowed the identification of 412 CDS. The genome has long direct terminal repeats (DTR) of 39,080 bases, a characteristic also found in previously published jumbo phages (Yoshikawa *et al.*, 2018). Interestingly, the DTR region includes 105 genes, many of which are highly transcribed in the early phase of the infection (Figure 2). Aside from 449 protein coding sequences, the phage genome also contains twelve tRNAs [Met (CAT), Leu (TAA), Asn (GTT), Thr (TGT), Met (CAT), Leu (TAA), Arg (TCT), Met (CAT), Pro (TGG), Gly (TCC), Ser (TGA) and Ser (GCT)] that were identified by tRNAscan-SE (Lowe et al., 2016) and confirmed in the RNA-Seq data. Transcriptomics data also revealed four actively transcribed regions lacking putative ORFs or known promoters that have been annotated as ncRNA (see Table 1).

**Table 1.** Identified non-coding RNA species. Four non-coding RNAs have been identified based on RNA-Seq data from transcripts aligning to genomic loci lacking plausible ORFs or known promoters.

| Name | Location | Strand | Percentage of phage reads |  |  |
| --- | --- | --- | --- | --- | --- |
|  |  |  | Early | Middle | Late |
| ncRNA001 | 8263..8498 | + | 2.40% | 5.07% | 4.68% |
| ncRNA002 | 21951..22401 | + | 4.65% | 5.04% | 1.23% |
| ncRNA003 | 103065..103368 | + | 0.01% | 0.05% | 0.03% |
| ncRNA004 | 246116..246734 | + | 0.08% | 0.03% | 0.02% |

**Figure 1.**
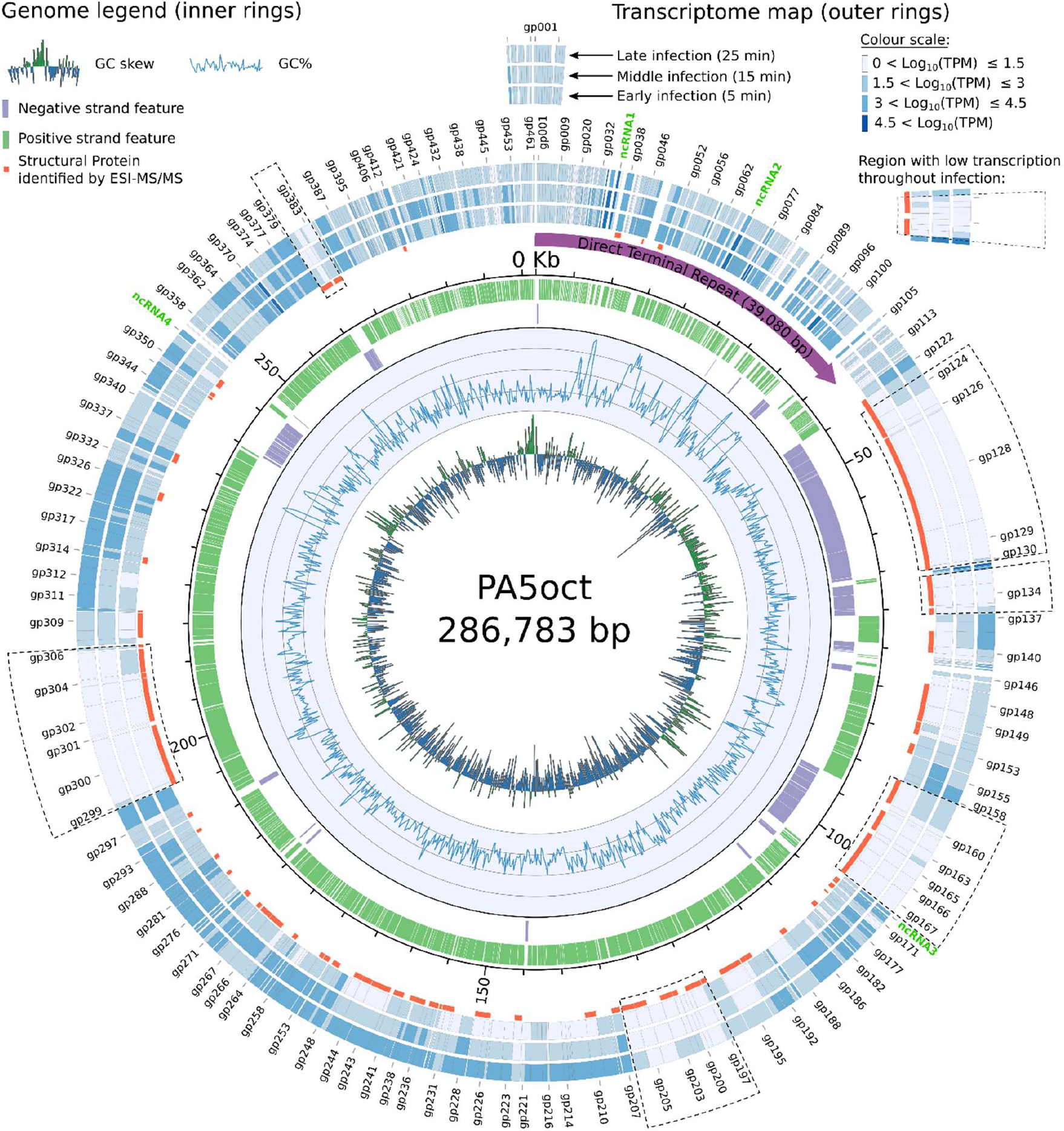
Circular representation of PA5oct genome and its transcriptome. The two inner circles display a histogram of the GC-skew followed by a graph of the GC content along the genome of PA5oct. It is followed by a stranded depiction of the CDS and ncRNA indicating that most features are encoded on the positive strand (in green) with intermittent regions encoding on the negative strand (in purple). Structural proteins confirmed by ESI-MS/MS analysis are highlighted in red color. The three outer rings depict normalized RNA read counts for each of the annotated feature (note that some gp numbers are not shown on the graph to ease the visualization). Contiguous regions with limited amounts of transcription activity detected across multiple timepoints are encased in dotted lines. The figure was created with the software *Circos* (Krzywinski *et al.*, 2009)

**Figure 2.**
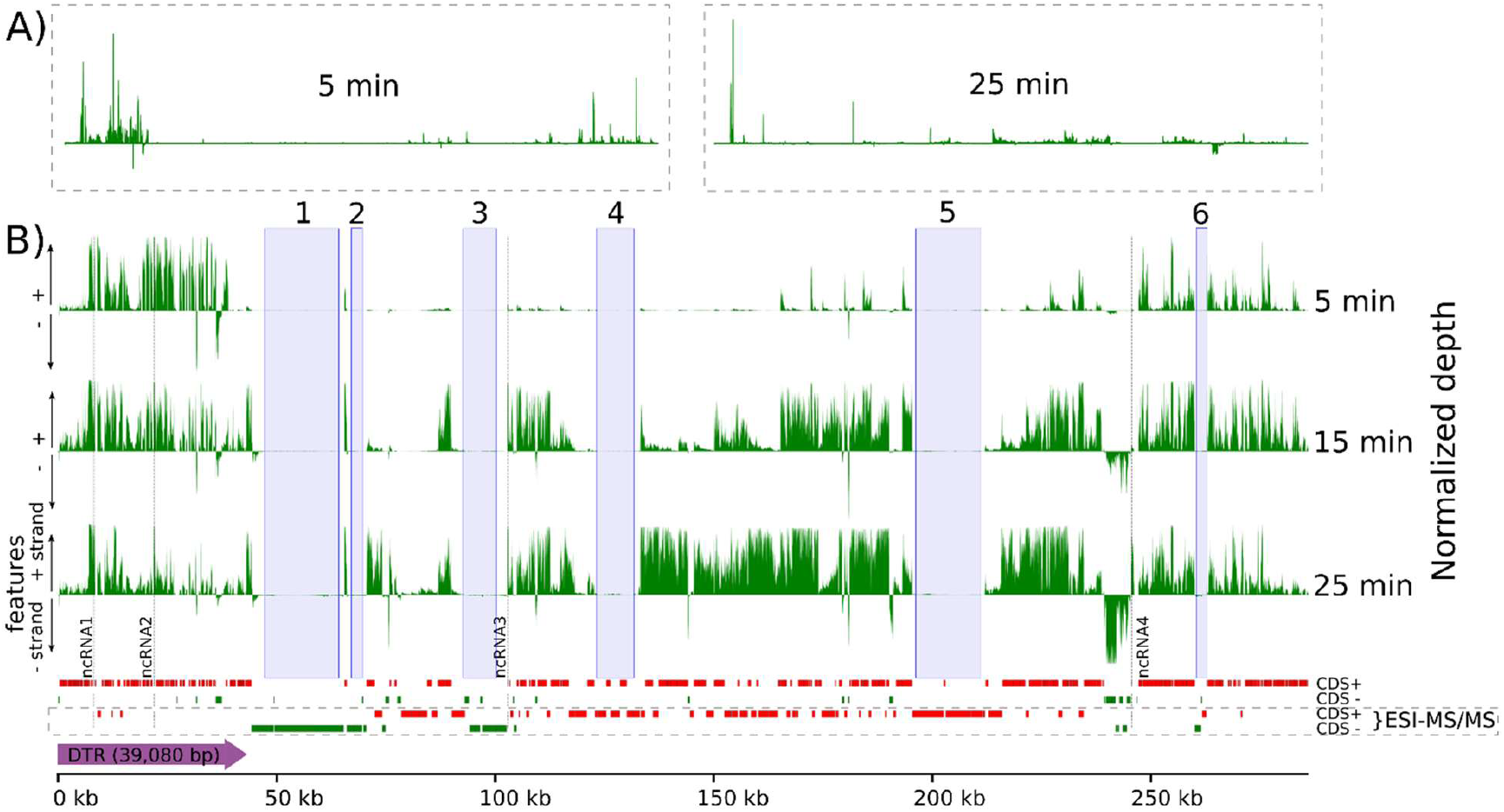
RNA-Seq analysis of PA5oct transcriptome after *P. aeruginosa* PAO1 infection. Genome-wide overview of reads mapped to the PA5oct genome for samples taken 5, 15, and 25 minutes post infection. A) Coverage of samples 5 min and 25 min (15 min omitted) show a limited number of regions which dominate the transcription landscape. B) Zooming in on the samples by capping the strongly transcribed regions allows to delineate regions differentially transcribed throughout the infection. Six marginally transcribed regions are highlighted with blue bars. Each of the three timepoints were normalized to reflect the depth across the triplicates. The figure was created using the python software *pyGenomeTracks*.

### Temporal regulation of PA5oct genome expression

PA5oct transcripts progressively come to dominate the host transcription. They represent 21% of total non-rRNA transcripts after the early infection timepoint (5 min), increase to 69% by middle infection (15 min), and culminate to 93% during late infection (25 min). The strand specificity of the results revealed that early transcription is almost exclusively occurring from the Watson strand.

Most of the genes contained within the long DTR between gp34 and gp105 (uncharacterized proteins, except gp39, 48, and 51, which are structural proteins), but also gp450 (uncharacterized protein), are each found to be highly transcribed at this stage and can be defined as early genes. Phage PA5oct appears to have few genomic regions that can be distinguished between the middle and late phase of transcription as captured at 15 minutes onwards (Figure 2-B). Many of the (functionally annotated) gene features transcribed in those middle and late timepoints are predicted to be responsible for structural proteins as well as DNA metabolism and replication. Two of the four non-coding RNAs found in the DTR region (ncRNA1 and ncRNA2), and which lack similarity to any known primary DNA sequence in the Genbank database (release 231), are highly transcribed throughout the infection cycle (Figure 1, 2 and Supplementary table S2).

Remarkably, six genomic regions, including two of the three major regions transcribed on the Crick strand (Figure 2-B, boxes 1-6), were found to show limited transcription under our laboratory growth conditions, despite a high sequencing depth (between 11M and 21M reads mapping for each individual sample). These six regions correspond primarily to genes that encode structural proteins (Figure 1, gene products marked in red) and which have been detected by ESI-MS/MS.

A total of 32 putative promoters with highly conserved, AT-rich intergenic motifs (5’-TATAATA-3’) and (5’-TTGAC-3’) were identified around transcription start sites (Supplementary table S2) whereas 36 putative factor-independent terminators with conserved stem loops could be identified around transcription stop sites (Supplementary table S3).

### Transcriptome and proteome-based functional annotations

The ESI-MS/MS technique allowed to delineate a total of 93 gene products linked to the phage particle, comprising experimentally confirmed seven virion-unrelated enzymes, 13 virion-associated proteins, and 73 other gene products (Figure 1S, supplementary table 2S). Combined with the bioinformatics analysis of the genome and the transcriptome, we could functionally annotate of a third of the CDS found in PA5oct.

Three capsid proteins were predicted, including the portal vertex protein of the head (gp149), the prohead core scaffolding protein and protease (gp152) and the precursor to the major head subunit (gp154). An additional protein, assigned as a putative head completion protein (gp200), was annotated based on sequence similarity to *Klebsiella* phage vB_KleM_RaK2 head completion protein. Moreover, several proteins associated with the tail apparatus were annotated, including a neck protein (gp193) and four tail proteins, a phage tail fiber protein H (gp159), a tail sheath stabilizer and completion protein (gp162), another tail fiber protein (gp165) and a tail sheath monomer (gp195) as well as three baseplate proteins, gp129, gp130 and gp133 (baseplate wedge, baseplate protein, baseplate hub subunit and tail lysozyme, respectively). Gp159, the putative phage tail fiber protein H, has a putative endo-N-acetylneuraminidase region on its C-terminus (868 – 1008 aa), suggesting a (2->8)-alpha-sialosyl linkage hydrolase function of oligo- or poly(sialic) acids, activity associated with the tail spikes and exopolysaccharide depolymerases (Kwiatkowski *et al.*, 1982).

PA5oct also encodes several genes predicted to be associated with peptidoglycan layer degradation, including three ESI-MS/MS detected product gp133, gp173, gp245, and two that were not detected gp368, and gp441. Their putative function was analyzed using BLASTP, HHpred, HMMER, InterPro, and Phyre2 (Table 2).

**Table 2.** Putative peptidoglycan degradation proteins. Five genes are predicted to be associated with the degradation of the peptidoglycan layer.

| Gp | ESI-MS/MS | Functional analysis |
| --- | --- | --- |
| gp133 | yes | Baseplate hub subunit with tail-associated lysozyme homologous to the gp5 of *Escherichia coli* bacteriophage T4 which has a needle-like structure attached to the end of the tail tube serving in the injection process (Kanamaru *et al.*, 2005). |
| gp173 | yes | Glycosyl hydrolase, lysozyme-like protein with a 159 amino acids long domain (11 - 170 aa). |
| gp245 | yes | Baseplate hub and tail muramoyl-pentapeptidase |
| gp368 | no | Cell-wall hydrolase domain protein with an N-acetylmuramyl-L-alanine amidase SleB domain cleaving the bond between N-acetylmuramoyl and L-amino acid residues. It is potentially an endolysin given that it was not recognized during ESI-MS/MS analysis |
| gp441 | no | Murein hydrolase, peptidase M23, with a 103 amino acids long domain (30 - 133 aa). |

### Gene-sharing relationships with other jumbo phages including PA5oct to other viruses

To represent genetic relationships between *Pseudomonas* phage PA5oct and other bacteriophages with genomes exceeding 200 kbp (Hendrix, 2009), a gene-sharing network was built. Viruses that do not show significant similarity to the jumbo were excluded for clarity. The resulting network was composed of 1,355 viral genomes (nodes) belonging to the *Myoviridae, Siphoviridae, Podoviridae*, or uncharacterized phages and 37,955 relationships (edges) between them.

As shown in Figure 3A, 24 jumbo phages were placed into the largest connected component (LCC) that predominates the double-stranded DNA phages, whereas 22 phages were placed into three isolated components. A majority (19) of the 24 jumbo phages in the LCC, which includes PA5oct, fall within a highly interconnected region (subnetwork) that is rich in the *Tevenvirinae* viruses (Upper right in Figure 3A; Table 5S). A subsequent analysis found 60 coherent groups of viral genomes (i.e., viral clusters, VCs) embedded in this subnetwork (Figure 3B; Table S5), of which six jumbo phages including PA5oct and *Escherichia* phages 121Q/PBECO 4, *Klebsiella* virus RaK2, *Klebsiella* virus K64-1, and *Cronobacter* virus GAP32 showed stronger connections to each other but weaker connections outside the group due to their more common genes (i.e., homologous protein clusters, PCs) than the rest of the network (Figure 3B). Notably, a further inspection of their relationships (phage K64-1 was excluded as it has an incomplete protein annotation profile in NCBI) uncovered its sub-structure where PA5oct shares less common PCs (∼11-15%) to the remaining members (Figure 3C), indicating its distant relationships as a outlier genome (Jang et al., 2019).

**Figure 3.**
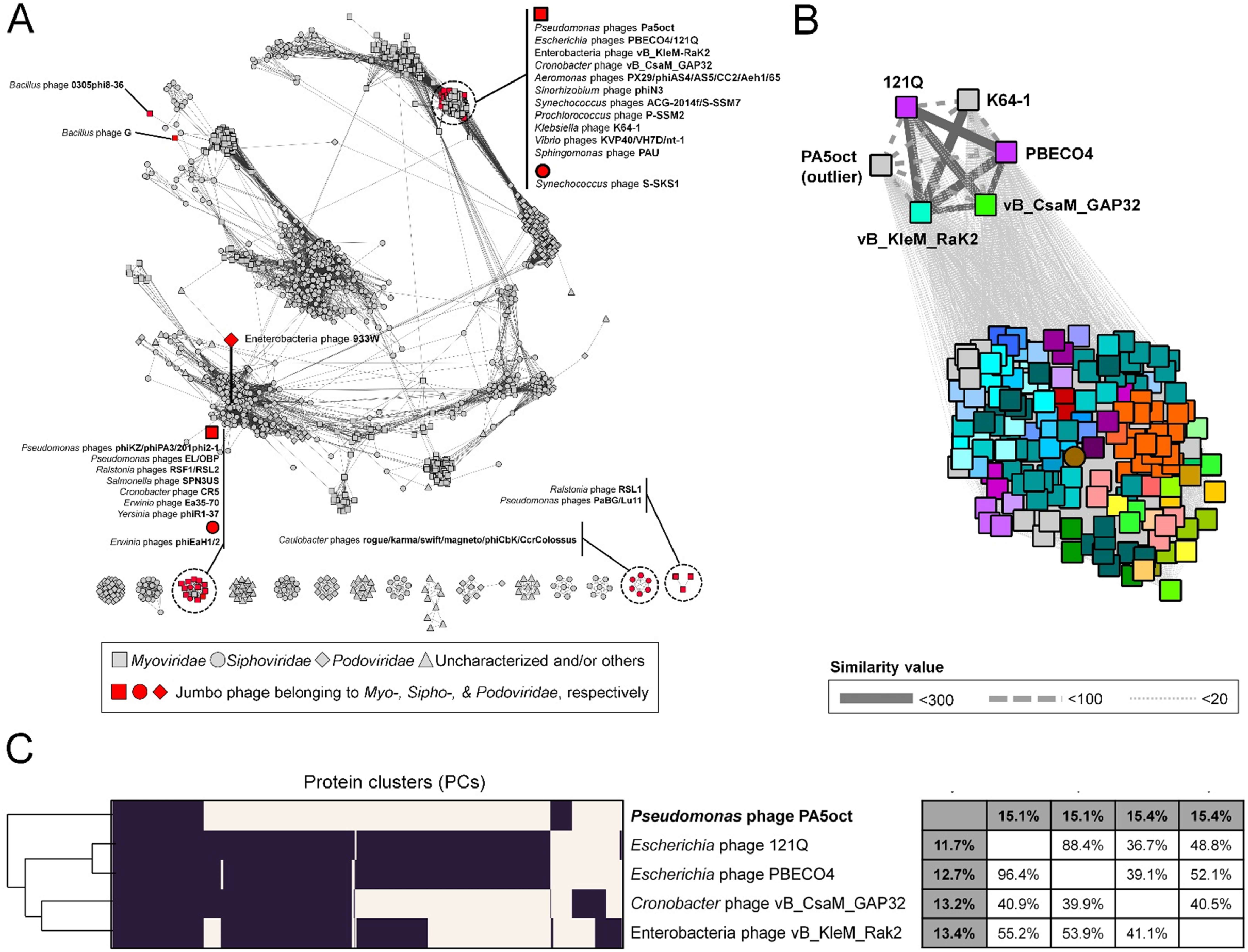
Protein-sharing network for PA5oct and other jumbo phages. (A) A network representation was produced using the edge-weighted spring embedded layout of Cytoscape version 3.5.1. Each node is depicted as a different shape, representing bacteriophages belonging to the *Myoviridae* (rectangle), *Podoviridae* (diamond), *Siphoviridae* (circle), or uncharacterized phages (triangle). Selected jumbo phages are shown in red. Edges between two nodes indicate statistically weighted pairwise similarities with similarity scores of ≥ 2. (B) An enlarged view of the subnetwork comprising PA5oct and its relatives. Edge thickness is proportional to similarity values estimated with the hypergeometric equation (Materials and methods) and viral clusters are denoted as different colors. Note that due to the lower number of common genes, *Pseudomonas* virus PA5oct was identified as a distant relative (outlier; Jang *et al*., 2019) to five phages including 121Q, K64-1, PBECO4, GAP32, and Rak2 (C) Module profiles showing the presence (dark) and absence (light) of homologous protein clusters (PCs) across genomes. Each row represents a virus and each column represents a PC. The genomes were hierarchically grouped based on hierarchical patterns of gene sharing (Left). A matrix showed the percentage of shared PCs between five-member viruses (Right). From these proteome-based analyses, phage K64-1 was excluded due to its incomplete protein annotation in NCBI.

With respect to other jumbo phages, the network identifies informative connections. For example, the *Phicbkvirus*, and phages RSL1, PaBG, and Lu11 form two isolated components (Fig. 3A). This discontinuous structure of two viral groups, due to their distinct gene pools, can reflect their evolutionary relationships as separate viral lineages (Gill *et al.*, 2012; Yamada *et al.*, 2010); whereas the connection of jumbo phages to their relatives having smaller genomes (<200kbp) such as *Bacillus* phage G, phages of the *Phikzvirus, Elvirus, Agrican357virus*, and *Rsl2virus*, and *Bacillus* phages 0305phi8-36Figure 3A and Table S5) can support evolutionary links between jumbo phages and smaller-genome phages (Hendrix, 2009; Adriaenssens *et al.*, 2012; Jang *et al.*, 2013). Together, the visualization and analysis of gene content relationships of the jumbo phages as a network revealed the global distribution of genetically diverse groups of jumbo phages across 2,259 viruses, in which most of them together with smaller-genome phage(s) are placed into two major groups comprising viruses belonging to the *Tevenvirinae* as well as the *Phikzvirus/Elvirus/Agrican357virus/Rsl2virus*, respectively, and others form evolutionary distinct clades (Gill *et al.*, 2012; Yamada *et al.*, 2010) (Table S5). In particular, despite the similarities of PA5oct to the recently characterized jumbo phages vB_CsaM_GAP32, 121Q, PBECO 4, vB_KleM-RaK2, and K64-1 (Abbasifar *et al.*, 2014; Ackermann *et al.*, 1983; Kim *et al.*, 2013; Šimoliūnas *et al.*, 2013; Pan *et al.*, 2015), follow-up analyses indicate that PA5oct is a more distant member of that group as evidenced by the fewer number of proteins shared with the group (Figure 3B and 3C).

### PA5oct-specific differential gene expression of the host genome

The conserved PAO1 host-mediated transcriptional response to phage infection present during the infection by phages 14-1, PEV2, YuA, and LUZ19 (Blasdel *et al.*, 2018) is also found during the lytic infection cycle of PA5oct (Figure 4). However, PA5oct also appears to elicit several phage-specific impacts on transcript abundance, which result in a unique differential expression pattern of specific host genes. PA5oct progressively dominates the non-ribosomal RNA environment of the cell and this shift from host transcripts to phage transcripts (Figure 4) depletes all host transcripts 0.08-fold relative to the total in the cell. Note that this global depletion of host transcripts relative to the total phage/host transcript population is not accounted for in the Log_2_ (Fold Change) values we report (Figure 4, Supplementary table S6), as described previously (Blasdel *et al.*, 2018). Doing this allows us to illustrate the impact of PA5oct on host transcripts relative to other host transcripts, rather than relative to the total.

**Figure 4.**
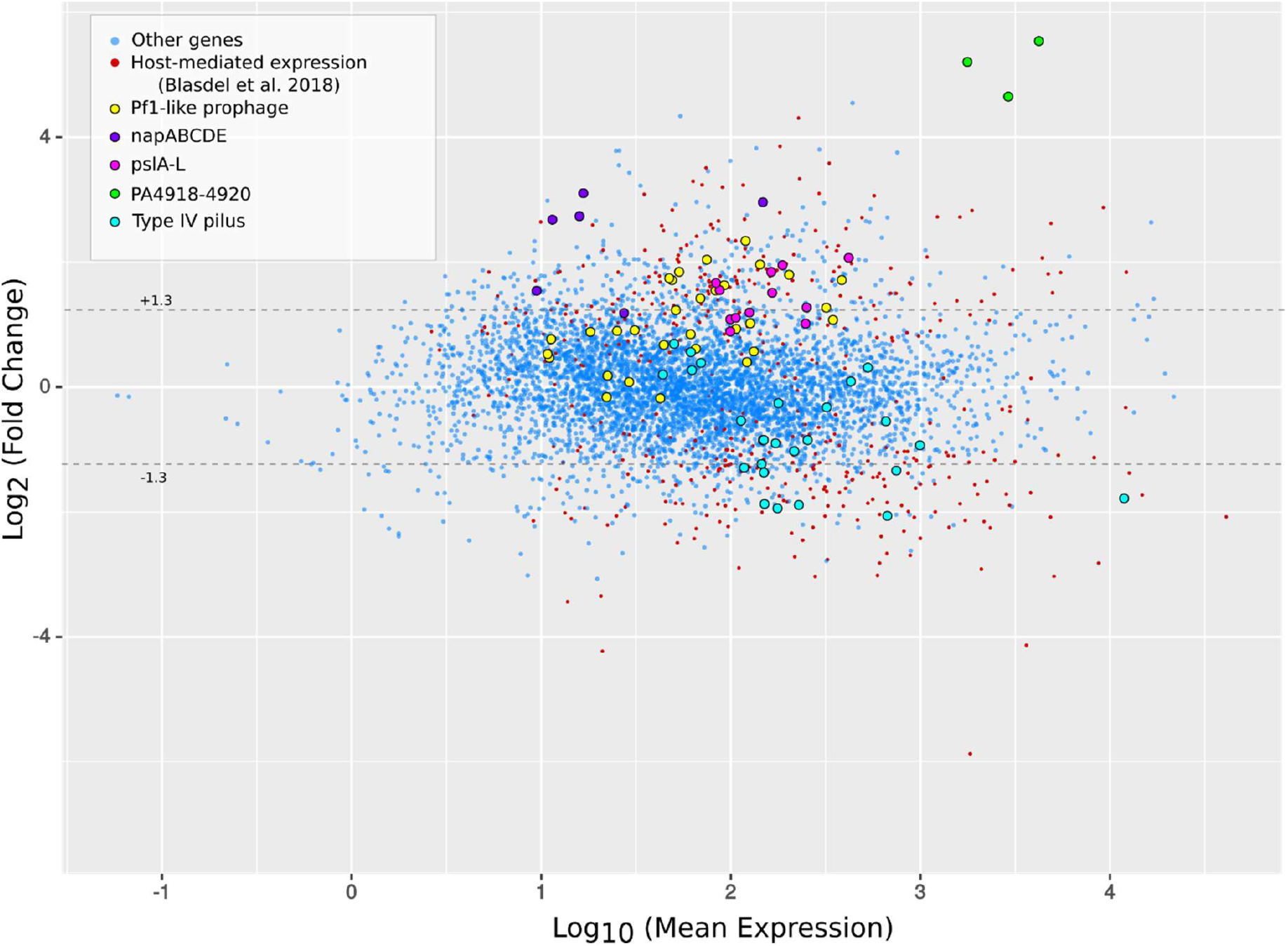
MA Plot highlighting the differential expression of host genes during PA5oct infection. In addition to the host-mediated stress response to phage infection identified by Blasdel *et al.* (2018), we observe PA5oct-specific differential expression of host genes during PA5oct infection. Specifically, we see a PA5oct-specific upregulation of the *pslE-J* part of the *pslA-L* exopolysaccharide operon, the *napABCDE* operon which encodes the periplasmic nitrate reductase, parts of a Pf1 prophage, and the operon PA4918-4920 involved in NAD biosynthesis. We also observe a downregulation of Type IV pili genes which is a necessary receptor to initiate PA5oct infection.

As shown on Figure 4, the strongest upregulation is that of the PA4918-PA4920 operon, which contains an enzyme involved in NAD biosynthesis (Okon et al., 2016). Another operon specifically upregulated during PA5oct infection encodes a part of the *psl A-L* gene cluster (*pslE-J)*, which is involved in the production of Psl exopolysaccharide (Jackson *et al.*, 2004). PA5oct also uniquely manipulates the host into upregulating the transcription of the *napABCDEF* operon, which encodes a periplasmic nitrate reductase. Interestingly, several genes associated with the biosynthesis of Type IV pili are downregulated relative to other host transcripts. As appears to be common in different infection settings with other virulent phages (Blasdel *et al.*, 2018), PA5oct appears to mobilize the transcription of genes of a potentially cryptic Pf1 prophage (family *Inoviridae*) encoded from ORF PA0618 to PA0642.

## Discussion

### Insights from the integration of multiple omics analyses

Phage PA5oct has currently the third largest genome amongst the sequenced *Pseudomonas* jumbo phages in available databases, after 201phi2-1 (316.674 kbp, Thomas *et al.*, 2008) and phiPA3 (309.208 kbp, Monson *et al.*, 2011). To date, the largest myovirus known is *Bacillus megaterium* phage G (497,513 bp genome, Table 1S). It is important to keep in mind that some jumbo phages, especially those reported prior to the 1990s, were identified only by electron microscopy (e.g. *Gluconobacter* phage GW6210) (Abbasifar *et al.*, 2014) and that ongoing sequencing efforts of these isolates, or metagenomics efforts will likely disrupt the current rankings of jumbo phage genome sizes.

In terms of annotation, functions could be attributed for a third of PA5oct proteins and a large portion of predicted CDS are unique, lacking sequence-based similarity to the genomes available in the public database Genbank (r231). For this reason, the functional characterization of jumbo phage orphan genes is an important endeavor to undertake in order to elucidate basic phage biology as well as phage application safety when thinking of therapeutic settings (Yuan *et al.*, 2017). Leveraging RNA-seq for a whole genome analysis is a powerful way to elucidate differential expression of gene features across different conditions but also delineate possible phage transcriptional modules. By experimentally defining the timing and expression levels of transcripts in both phage and host, directional RNA-Seq helps to discover and accurately annotate novel sequences, particularly for ncRNAs that are difficult to predict using genomics data only, but also for small phage peptides that fall below gene prediction thresholds of annotation pipelines (Ceyssens *et al.*, 2014). In this manner, we could refine annotations of existing coding sequences, correcting and adding about 10% of the total predicted ORFs from those predicted *in silico* as well as annotating four ncRNA regions of high interest for future investigation. Interestingly, we also discovered a ribonuclease H-like protein on the genome of PA5oct (gp325), putatively involved in the processing of the host transcripts during the infection cycle.

ESI-MS/MS allowed us to identify 93 structural proteins of PA5oct, including four structural head related proteins, one neck protein and four tail related proteins. This is a similar result to that seen with giant *Pseudomonas* 201phi2-1 (89 structural proteins, Thomas *et al.*, 2010). However, other jumbo phages can possess a much smaller number of structural proteins, such as *Pseudomonas* virus phiKZ (62), *Aeromonas* virus phiAS5 (26) or *Ralstonia* virus phiRSL1 (25) (Lecoutere *et al.*, 2009; Yamada *et al.*, 2010; Kim *et al.*, 2012). Furthermore, the capsid of jumbo phages can have very complex structures, as we could observe in the case of phage phiKZ (30 head structural proteins), where in the middle it has an “inner body”, a spool-like protein structure that plays an important role in DNA packaging and genome ejection during, respectively, phage virion assembly and subsequent virion adsorption (Wu *et al.*, 2012).

Importantly, the large capsids seen in jumbo phages can accommodate large genomes allowing these phages to possess many accessory genes comparing to small phages (Ceyssens *et al.*, 2014; Yuan & Gao, 2017; Mesyanzhinov *et al.*, 2002; Hertveldt *et al.*, 2005; Kiljunen *et al.*, 2005; Thomas *et al.*, 2007; Gill *et al.*, 2012). For example, PA5oct encodes five different genes encoding peptidoglycan-degrading enzymes and 12 tRNA genes to increase the translation efficiency (Yuan & Gao, 2016). Note that other jumbo phages may contain even more tRNAs genes, such as *Vibrio* virus phi-pp2 and KVP40 (30 tRNA); *Caulobacter* virus CcrColossus and *Vibrio* virus nt-1 (28 tRNA); *Aeromonas* virus Aeh1, *Caulobacter* virus Swift and Magneto (27 tRNA); *Cronobacter* virus GAP32, *Caulobacter* virus Karma and phiCbK (26 tRNA); *Aeromonas* virus PX29 (25 codons) (Supplementary table 1S).

### Hijacking the transcriptional environment of the host

The strategy of phage transcriptional mechanisms depends primarily on the presence or absence of a phage RNA polymerase (Yang *et al.*, 2014). After careful analysis of PA5oct genes, no RNAP-like proteins could be identified. Taken together with the absence of phage production in the presence of rifampicin and the host’s ability to mount a transcription-level defense response, this suggests that this phage may depend on host transcriptional mechanism, though we cannot rule out the possibility that an unknown but essential phage protein is also sensitive to rifampicin. The main role in phage gene transcription is thus likely played by the large DNA-dependent RNAP of bacterial host, a 400-kDa protein complex composed of five subunits (α2, β, β’,ω), that transcribes mRNA following binding to the DNA template (Murakami, 2015; Tagami *et al.*, 2014, Davis *et al.*, 2017). This process is often supported by phage-encoded σ factors forming a holoenzyme with RNAP (α2, β, β’,ω, σ) (Williams *et al.*, 1987; Pavlova *et al.*, 2012).

Some phages can also implement other transcriptional strategies, e.g. viral gene transcription may rely on both the host RNAP along with a single-subunit phage RNAP (*Enterobacteria* virus T7, T3, *Pseudomonas* virus phiKMV and *Xanthomonas oryzae* virus Xp10). The single-subunit RNAP is then responsible for transcription of phage genes in the middle and late stage of infection, whereas early genes are transcribed by the host RNAP (Savalia *et al.*, 2010; Semenova *et al.*, 2005, Yang *et al.*, 2014). A similar situation is observed in the case of *Enterobacteria* virus N4, where early and middle stage of transcription depends on two phage-encoded RNAPs and the late genes are transcribed by host RNAP (Haynes *et al.*, 1985; Willis *et al.*, 2002). Another example is the giant *Pseudomonas* virus phiKZ that can infect its host independently of the host RNAP by using a virion-associated phage RNAP active in early transcription, and a non-virion-associated phage RNAP, important for late and possibly middle transcripts (Ceyssens *et al.*, 2014; Yakunina *et al.*, 2015).

### Phage transcription patterns

PA5oct progressively dominates host transcription. After 5 min, PA5oct transcripts represented 21% of total non-rRNA transcripts, eventually proceeding to 69% and then 92% by middle and late infection. This possibly reflects a globally accelerated degradation of RNA in the cell akin to what was described by Chevallereau *et al.* (2016). In a similar fashion to giant *Pseudomonas* virus phiKZ (Ceyssens *et al.*, 2014), giant Yersinia virus phiR1-37 (Leskinen *et al.*, 2016) and giant *Bacillus* virus AR9 (Lavysh *et al.*, 2017), PA5oct is not organized into strictly contiguous regions of temporal transcription. In the early timepoint, we observe a lot of transcription in the DTR region (Figure 2). These early coding sequences are classically understood to be involved in the transitioning from host to phage metabolism. Other regions of early, middle, and late expression are scattered throughout the genomes, a feature that increasingly appears to be characteristic of phages with large genomes, including the 168 kbp *Escherichia* virus T4.

We also found six regions with low levels of transcription that encode for unknown structural genes correlated to baseplate and tail that were confirmed to be translated and incorporated into phage particles by ESI-MS/MS (a very sensitive technique), suggesting either a notably efficient translation or a novel and unknown bias during RNA-seq. It is tempting to speculate that the expression of these structural proteins may be conditional, for example based on environmental cues which result in altered tail-receptor proteins to be displayed. Indeed, these specific structural proteins are not conserved between phages within the protein-sharing network, hinting at functions associated with specificity. These genes could have been captured from other phages and be part of the accessory genetic content of these jumbo phages.

### Comparative genome analysis and protein-sharing network

To examine the genetic relationships between PA5oct and other jumbo bacteriophages as well as *Myoviridae, Siphoviridae, Podoviridae* viruses, a protein-sharing network was constructed. The PA5oct, *Escherichia* viruss 121Q/PBECO4, *Klebsiella* virus vB_KleM-RaK2, *Klebsiella* virus K64-1, and *Cronobacter* virus vB_CsaM_GAP32 presented closer relationships based on shared conserved core genes. Furthermore, PA5oct, PBECO4, GAP32, T5, FelixO1 appear to be distantly diverged members of *Tevenvirinae*. The T5 and FelixO1 phages appear also to act as bridge for T4-related components due to their links to other phage groups. Unlike the marker genes of bacteria (i.e., 16S rRNA genes), which can be used for their taxonomic classification, viruses lack universal genes. Thus, traditional single-gene-based phylogenies, such as those based on major capsid proteins, will always be limited to related subsets of the total virus universe. Protein-sharing networks, as presented here, introduce an alternative approach, which provides a more global view that is also consistent with current taxonomic groupings. Using this approach, the possible genetic relationships for PA5oct, but also all giant virus among the whole virus population can be observed.

### Host stress response during PA5oct virus infection

Phages have a substantial impact on bacterial cellular systems, including transport/export, energy production/conversion, ribosomal proteins, cell wall modification, conversion of ribonucleotides to deoxyribonucleotides, cold shock and osmotic stress response (Fallico *et al.*, 2011; Ainsworth *et al.*, 2013; Ravantti *et al.*, 2008; Leskinen *et al.*, 2016). In this study, we observed that PA5oct upregulates the operon encoding *psIE-J*, responsible for production of Psl exopolysaccharide. Psl is essential for initiating and maintaining biofilm architecture in both mucoid and non-mucoid strains. It is responsible for the formation of a fabric-like matrix that holds cells closely together, thereby impeding migration within the biofilm (Ma *et al.*, 2009). This may lead to increased slime production, reducing the relative abundance of LPS phage receptors, hence secondary phage infections.

The *napABCDEF* operon, involved in the production of a periplasmic nitrate reductase, is upregulated during PA5oct infection. This protein is required for anaerobic growth in cystic fibrosis (CF) sputum *in vitro*. In a CF environment, *P. aeruginosa* prefers anaerobic growth that significantly enhances its biofilm formation and antibiotic resistance (Yoon *et al.*, 2002). Under these anaerobic conditions, *P. aeruginosa* modifies the structure of its LPS from a highly electronegative surface to a neutral surface (Sabra *et al.*, 2003, Palmer *et al.*, 2007). This suggests that phage PA5oct may influence *Pseudomonas* biofilm formation. While the purpose and mechanistic origin of this upregulation is unclear, it would seem to suggest that PA5oct may be well adapted to the environment of human-associated *Pseudomonas* biofilms, an interesting prospect for phage therapy given this phage’s broad host-range.

The upregulation of the operon PA4918-4920 can be linked to NAD biosynthesis, an important molecule that mediates the transfer of energy in the cell, potentially part of the phage mediated conversion of the bacterial cell into an optimized phage production system.

We observed here again an upregulation of prophage pf1, a key filamentous prophage that is linked to the pathogenicity of *P. aeruginosa* in lung infections (Burgener *et al.*, 2019), which appears to be ubiquitous during phage infection. Indeed, this has been observed in other transcriptomics assays of phage infection with phiKZ (Ceyssens *et al.*, 2014), as well as with phages LUZ19, PEV2, PakP3, and PakP4 (Blasdel *et al.*, 2018). It can reasonably be proposed to be the result of a general stress response of phage Pf1 that is unrelated to superinfection exclusion mechanisms.

Since PA5oct downregulates biosynthesis of Type IV pili and indirectly modifies bacterial LPS, the signaling networks might also influence the overproduction of nitrate reductase. As it was proven in our related manuscript, Type IV pili and LPS structures serve as phage PA5oct receptors (Olszak *et al.*, 2019). Receptor downregulation possibly functions as a tactic to reduce losses of “sister” phages and progeny adsorption (i.e., secondary adsorption) to already phage-infected bacteria (Abedon, 2017).

## Funding

This study was supported by research grant no. 2012/04/M/NZ6/00335 of the National Science Centre, Poland. KDW was co-financed by the European Union as part of the European Social Fund and a postdoctoral fellowship at KU Leuven, Belgium. CL is supported by an SB PhD fellowship from FWO Vlaanderen (1S64718N). RL, YB, and ZDK are members of the “PhageBiotics” research community, supported by the FWO Vlaanderen. ZDK was a members of the EU COST BM1003 Action in Microbial cell surface determinants of virulence (http://www.cost.eu/COST_Actions/bmbs/BM1003). JPN is supported by a grant from the Hercules Foundation (project R-3986).

## Acknowledgements

We are grateful to Steve Abedon and Harald Brüssow for critical comments and helpful suggestions on the manuscript.

## Authors contributions

KDW performed one-step growth experiments and analysis of ESI-MS/MS results. CL, KDW performed DNA isolation, and annotation of the genome. HBJ generated protein family clustering, network construction and analyzed the dataset. CL, DV helped with promoter identification. CL, KDW, YB performed the genome sequencing and assembly. JPN performed ESI-MS/MS analysis. BB and KDW prepared samples for RNA-Seq. BB, CL, KDW analyzed the transcriptomics results. BB, CL, HBJ, KDW, RL, ZDK designed experiments and wrote the manuscript.

## Supporting Information

**Supplementary table 1S. List of jumbo phages found in the litera**ture.

S**upplementary table 2S. Alignments of PA5oct promoters.** Conserved AT-rich intergenic motifs: −10 box (5’- TATAATA - 3’) and −35 box (5’ - TTGAC - 3’). The corresponding sequence logos from MEME/MAST are depicted below the alignments. ORFs in front of which promoters are located are listed on the left side, whereas the corresponding sequence locus in the phage genome is indicated on the right side.

**Supplementary table 3S. Phage PA5oct predicted terminators with palindromes marked blue.** From the left site: list of ORFs after which terminators are located by ARNold with strandedness, location in phage genome, motif of terminator, terminator sequence with conserved stem-loop structures marked blue.

**Supplementary table 4S. The ESI-MS/MS analysis of denatured phage PA5oct particles after fractionation on SDS-PAGE gel.**

**Supplementary table 5S. List of jumbo phages (yellow color) and relevant phages that belong to the viral clusters (VCs) and their ICTV taxonom**y.

**Supplementary table 6S. The differential expression of non-rRNA gene feature within *Pseudomonas* PAO1 genome by late PA5oct infection.**

**Supplementary figure 1S. The SDS-PAGE pattern of PA5oct structural proteome against LMW Ladder (Thermo Scientific) in the first lane.** The corresponding molecular weight is mentioned left. The numbered fractions on the right, correspond to gel slices analyzed individually by ESI-MS/MS. The proteins are mentioned in the slice in which they were most abundantly present (see Band no. in Supplementary Table S4).

