## Supplementary figures and images for "Integrative omics analysis of *Pseudomonas aeruginosa* virus PA5oct highlights the molecular complexity of jumbo phages"

### Supplemental figure S1

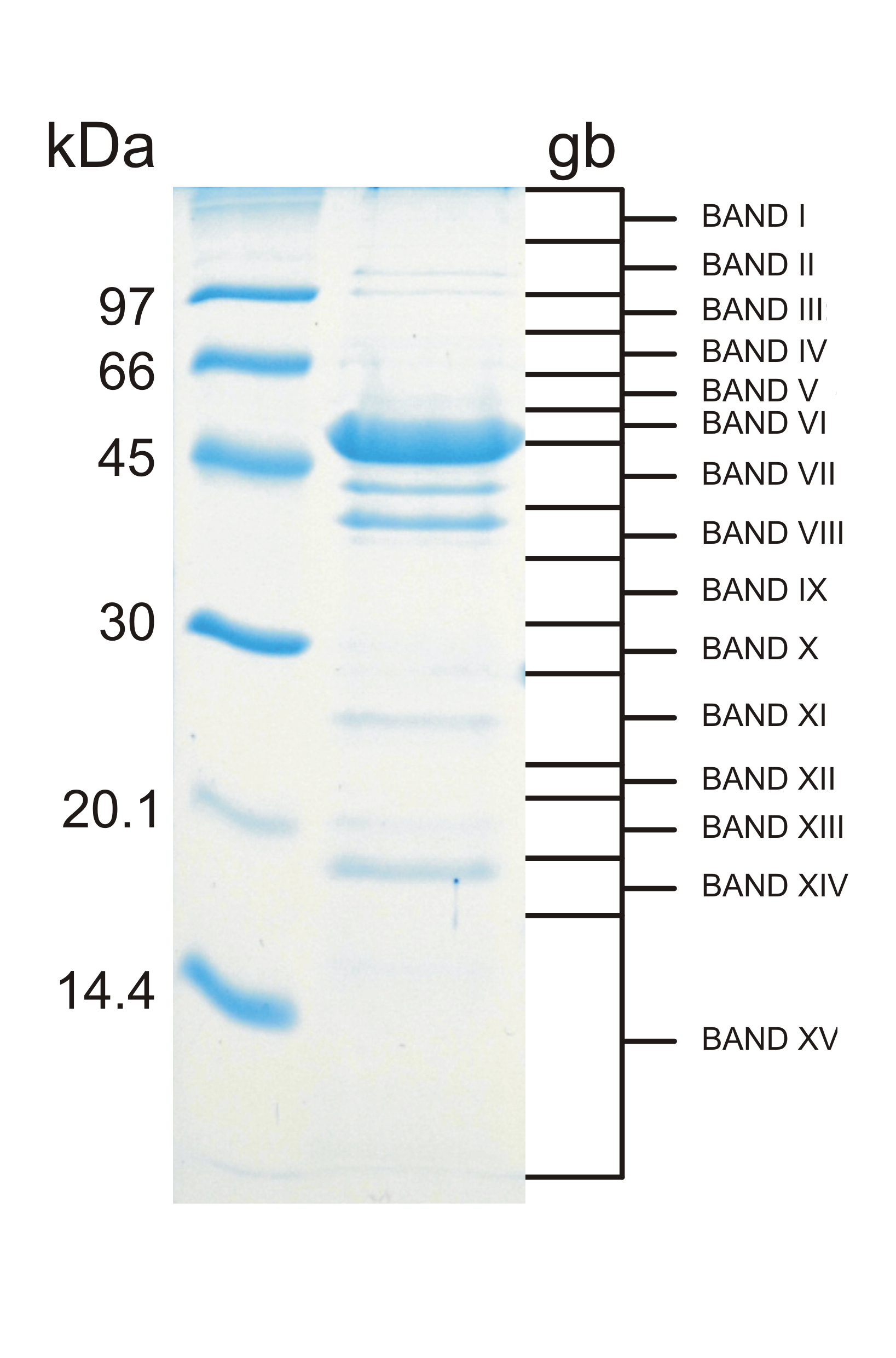
